# Dataset-specific thresholds significantly improve detection of low transcribed regulatory genes in polysome profiling experiments

**DOI:** 10.1101/2020.12.14.422662

**Authors:** Igor V. Deyneko, Orkhan N. Mustafaev, Alexander А. Tyurin, Ksenya V. Zhukova, Irina V. Goldenkova-Pavlova

## Abstract

**Motivation:** Polysome profiling is novel, and yet has proved to be an effective approach to detect mRNAs with differential ribosomal load and explore the regulatory mechanisms driving efficient translation. Genes encoding regulatory proteins, having a great influence of the organism, usually reveal moderate to low transcriptional levels, compared, for example, to genes of house-keeping machinery. This complicates the reliable detection of such genes in the presence of technical and/or biological noise.

**Results:** In this work we investigate how cleaning of polysome profiling data on *Arabidopsis thaliana* influences the ability to detect genes with low level of total mRNA, but with a highly differential ribosomal load, i.e. genes translationally active. Suggested data modelling approach to identify a background level of mRNA counts individually for each dataset, shows higher power in detection of low transcribed genes, compared to the use of thresholds for the minimal required mRNA counts or the use of raw data. The significant increase in detected number of regulation–related genes was demonstrated. The described approach is applicable to a wide variety of RNA-seq data. All identified and classified mRNAs with high and low translation status are made available in supplementary material.

## 1. Introduction

Investigation of the mechanisms underlying differential gene expression is one of the fundamental tasks in understanding the functional organization of genomes and their dynamic properties. To date, most attention has been focused on the stage of transcriptional regulation, partly due to the relative simplicity and the variety of established experimental techniques. From another side, there is a growing number of studies showing a large discrepancy between levels of transcription and the levels of the target proteins, suggesting the importance of the intermediate steps like the regulation of translation (also called ‘translational buffering’) [1–3]. One of the most fascinating studies shows that fluctuations in transcriptomes do not necessarily lead to changes in the protein levels [1]. This discrepancy is mainly attributed to the active regulation of translation. The rise of novel experimental techniques such as polysome profiling and ribosome profiling [4] forms a solid ground for deciphering such regulation. The basic idea behind all of these techniques is to separate mRNA in a quiet state (monosomal fraction) and active state, i.e. mRNA heavily loaded with ribosomes (polysomal fraction), followed by sequencing or hybridizing on chips [3]. The resulting quantitative measure of translational state allows a better correlation of the number of mRNA transcripts and the observed protein levels [5]. Additionally, such data can be used to investigate regulatory mechanisms of the observed differential translation.

There are a number of programs used for analysis of ribosome sequencing data, most of which were originally developed for the analysis of gene transcription [6–8]. The major problem of the mathematical methods behind these programs is the estimation of the variance, that is the key point for the calculation of the statistical significance of the observed differences. Estimation of the variance of the measured expression values can be based on variations between replicates or in more advanced approaches, on genes from the same replicate with similar absolute expression [7]. This allows having even a single sample to estimate gene expression variance and then a statistical significance of differences between genes.

Some programs were specifically developed for analysis of polysome and ribosome profiling experiments, which are usually designed to measure polysomal and total mRNA fractions. Programs like anota2seq [9] or RiboDiff [10] can directly adjust their mathematical models for the changes in total level of transcription. The idea behind anota2seq is to pool genes with similar transcription to increase statistical power using the generalization of random variance model [11], when the number of replicates is not sufficient.

Still, there are other factors, apart from variability, affecting statistical calculations, such as outliers and noise, that cannot be fully considered by these programs. The problem of removing the noise and the selection of the “correct” threshold for minimal value of mRNA count is very controversial, and there is no agreement on this in the bioinformatics community. In anota2seq [9] RNA counts equal to zero are automatically removed. DESeq2 [7] performs independent filtering by default using the mean of normalized counts as filter statistics. Software Corset [12] filters any transcripts with fewer than ten reads by default and in the analysis of microRNAs, it was suggested to set the threshold to 32 reads [13].

In this work it is suggested to define a threshold for the minimal required mRNA count based on the analysis of the investigated datasets. We demonstrate that this approach is more effective, compared to universal, pre-defined thresholds, especially in searching genes with low transcription, *i.e.* with low values of the measured mRNA counts. This approach can also be used for the analysis of transcriptome RNA-seq data and the idea of data modelling can be applied to any suitable dataset.

## 2. Materials and Methods

### 2.1. Plant material

Plants of *A. thaliana* type Columbia-0 were grown at 22°C, 12h lighting period, light intensity of 100 μmol*m-2*s-1 and sampled on the stage of third rosette leaf (approx. 28 days). Three independent samples were prepared.

### 2.2. Preparation of monosomal, polysomal and total mRNA fractions

Plant material (leaves) was homogenized in a buffer containing 0.2 M Tris pH 9.0, 0.2 M KCl, 0.025 M EGTA, 0.035 M MgCl2, 1% DOC, 1% Triton, 5 mM DTT, 50 mg/ml cycloheximide, 50 mg/ml chloramphenicol. Cell extracts were applied over 5 ml of a 15-60% (W/v) sucrose gradient and centrifuged at 237000g for 1.5 hours at 4 ° C. Fractions with a volume of 400 μl were taken manually. Total RNA was extracted from each fraction using the ExtractRNA kit (Evrogen, Russia). In each fraction, the RNA content was evaluated using a Nanodrop ND-1000 instrument (LabTech International, UK).

Total cytosolic RNA was isolated from the part of the cell extract before loading onto the sucrose gradient. RNA was extracted using the ExtractRNA kit (Evrogen, Russia), the quality and quantity of preparations of total RNA and RNA from polysomal and monosomal fractions of plants was evaluated on an Agilent Bioanalyzer 2100. More detailed description of the protocol can be found in [14]. Altogether, nine samples were prepared for sequencing.

### 2.3. Preparation of RNA samples, sequencing, assembling and mapping

RNA libraries were prepared with TruSeq Stranded mRNA Sample Prep Kit (Illumina), quality control were performed on Agilent Bioanalyzer 2100 and by qRCR. Sequencing was done on Illumina HiSeq 4000 (101 cycle, paired end) with HiSeq 4000 sequencing kit version 1. FASTQ files were filtered to remove adapters, low-quality reads and reads with more than 10% mismatches.

### 2.4. Statistical analysis

All statistical calculations were done in R [15] and MS Excel. Statistical difference between polysomal and monosomal fractions were calculated using edgeR version 3.24.3 with default arguments [6]. Fitting the exponential model was done using lm(log(#mRNAs)~mRNA_count) function in R. Differences in functional classifications are evaluated using binomial test. Genomic sequences were downloaded from EnsemblPlants (http://plants.ensembl.org/index.html) and processed using Perl scripts. Gene ontology analysis was performed using DAVID [16] and PANTHER v.14.0 [17].

## 3. Results and Discussion

### 3.1 Polysome profiling experiment

Protein production is a multistep process including transcription, transport, mRNA maturation, translation and final protein modifications. One way to study the regulation of translation is to measure the differential ribosomal load by polysome profiling [4]. Briefly, the method consists in mRNA extraction, separation in sucrose gradient into mRNA fractions with high (polysomal fraction) and low (monosomal fraction) ribosomal load [18]. mRNA released from ribosomes is sequenced, reads are mapped to the genome, count values for mRNA are calculated and analyzed with programs like DESeq2 or edgeR [6, 7], designed for differential analysis of NGS data and available as R [15] packages.

In this work, in addition to classical polysome profiling experiment design, the measurement of total cytosolic mRNA was also included. It was based on considerations, that mechanisms of translational regulation may be different in classes of abundant and rare mRNAs. Indeed, the regulation of rare mRNA is thought to be very sensitive, as for example, for genes encoding regulatory factors, where from a few mRNA copies many protein molecules can be produced via intensive translation. Taking into account the possible variety of the gene regulatory mechanisms on stages of transcription and translation, it seems necessary to be able to isolate groups of mRNAs similar not only by translational status, but also by transcriptional. Altogether, our experiment consists of measuring the levels of mRNAs in polysome, monosome and total cytosolic mRNA fractions, each performed in three replicates (Figure 1).

**Figure 1.**
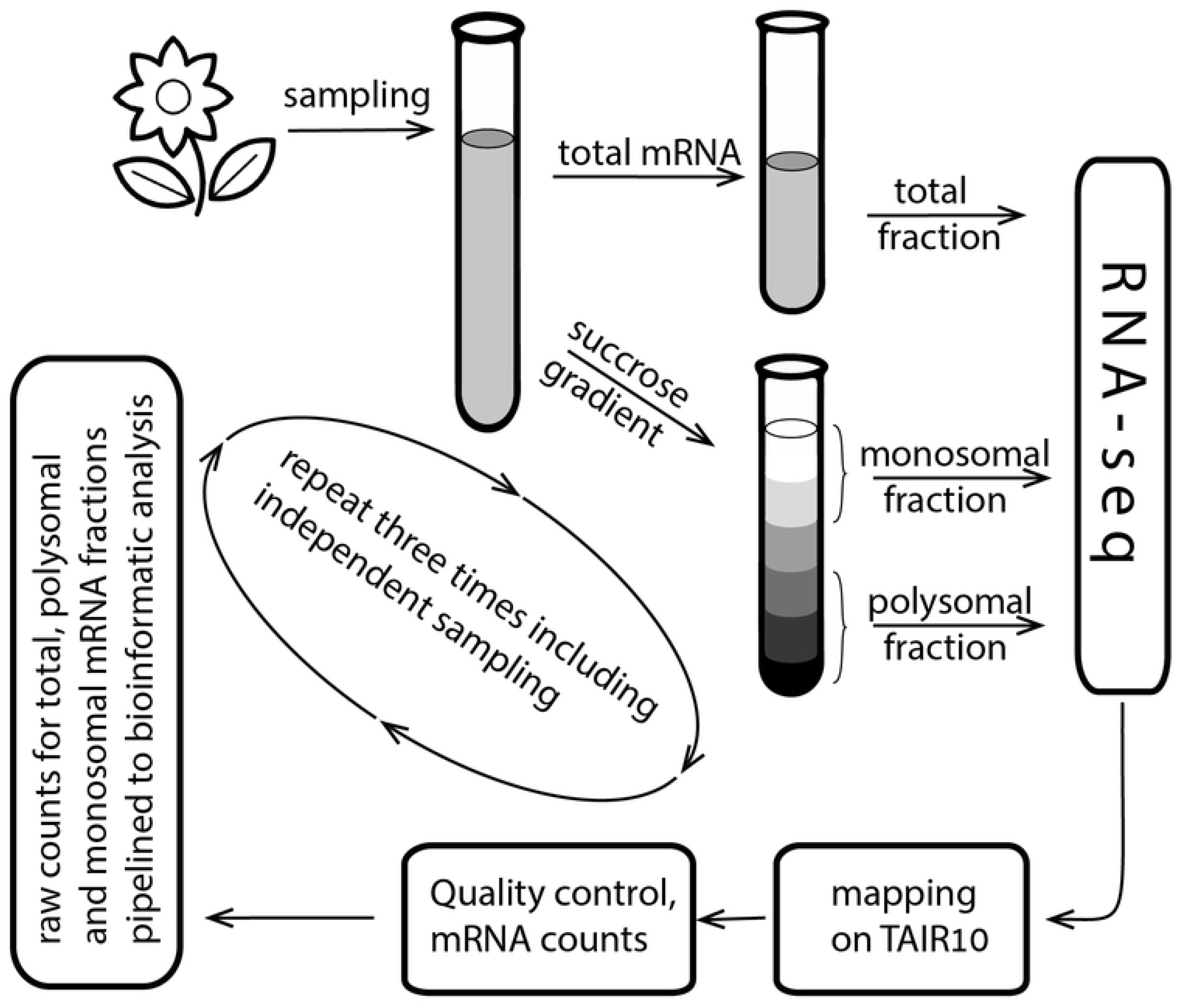
Schematic representation of the experimental design.

### 3.2 Modelling the raw data

Raw RNA counts coming from sequencing represent the amount of RNA found in the sample. In total 610M reads and 89G bases were sequenced, which were mapped to 37336 different mRNAs on the TAIR10 genome. Let *N_f,i_* be the number of reads for mRNA *i* = 1,…, 37336 in fraction *f=(polysome, monosome, total)*, averaged over the three replicates. Figure 2 represents the number of mRNAs with respect to their counts (*N_f,i_)*. It is interesting to observe a very high number of mRNAs with close to one counts, which decays as count number increases. Usually these small counts are regarded as noise and mRNAs with counts less than some predefined values are removed [7, 12, 13]. Here we suggest modelling the data distributions and to find exact values which should be subtracted from the raw values.

**Figure 2.**
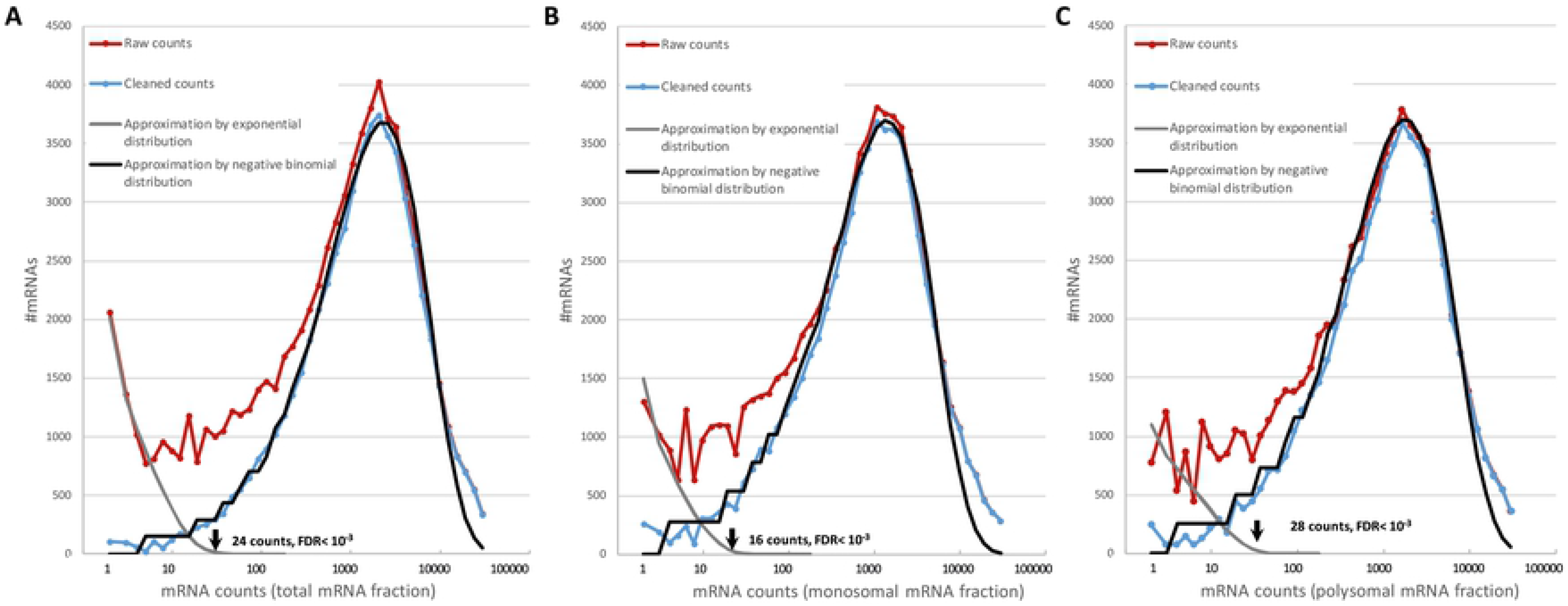
Distribution of mRNAs according to mRNA counts. These graphs show how many mRNAs have specified number of counts (empirical distributions, red curves) and its approximation by the exponent in the area of low values (grey dashed curves). Data, cleaned by subtraction the specified count value from every mRNA, is shown by the blue curves. The cleaned data is very close to negative binomial distribution (black curves). Graphs represent A) total B) monosomal C) polysomal mRNA fractions.

Overall, the distributions have two local maxima – one is around one and the other is around 3400 counts for total RNA fraction (2800 and 2500 for monosomal and polysomal fractions). One can speculate that this curve represents a sum of two independent processes, one is exponentially distributed and the other distributed negative binomially. The former can be interpreted as a background noise, which usually decay exponentially [19], and may originate from DNA debris, reverse transcription or sequencing artefacts. The letter is a real signal that has negative binomial distribution [20]. Formally this can be represented as a sum of two independent random variables, one following negative binomial distribution and the other exponential:

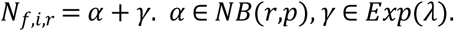

In other words, it is assumed that every measured mRNA count value contains real and random parts. It is not possible to decompose each value of mRNA count into two components due to the random nature of the process, but one can estimate the maximum contribution of the exponential part and then subtract it from the raw value. It is possible, because the contribution of the binomial part with its peak around 3000 is negligible at low values, therefore it will be assumed that points with very low values are of pure random nature.

The exponent distribution has one parameter and can be found by fitting the exponential model into data below ten counts (first several points on the red curve, fig. 2). Having built the exponential model (grey dashed curve, fig.2), one can extrapolate the curve to the point where the exponent drops to some acceptably low value, or in other words, solve for *m* the equation e^−αm^=10^−3^, where α is the estimated decay parameter. For example, the exponent equals 10^−3^ when mRNA count equals 24 for total mRNA fraction. That means, that one mRNA out of thousand with the count value of 24 is expected to appear by chance. The value of 24 can be used as a threshold for the minimal required counts instead of pre-defined threshold [7, 12, 13]. But following our logic, that the observed counts consist of two independent components, this value should be subtracted from all raw mRNA count values to maximally exclude possible random effect. If the resulting value is negative, a zero value is assigned:

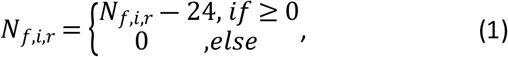

The distribution of the cleaned data is now very close to negative binomial distribution as it is usually assumed [6, 21] (blue curves, fig. 2). Overall, the three datasets of total, monosomal and polysomal fractions were modified by subtracting 24, 16 and 28 from each mRNA count respectively. So for example, if mRNA for a transmembrane protein gene AT3G55790 has 95 raw counts in first repetition of total mRNA fraction, then 95-24=71 counts will be the cleaned count value for that gene. After cleaning, mRNAs with all zero counts were removed, resulting in 23102 mRNAs out of 37336 in the raw data.

Evidently, this transformation mainly affects mRNAs with low counts and have no or minor effect on highly transcribed mRNAs. In the next section, the advantage of data-specific thresholds and the suggested data modification will be shown for detection of genes with regulatory function.

### 3.3 Detection of signal transduction and regulatory related genes is sensitive to the data cleaning procedure

Genes encoding regulatory proteins, including so-called master regulator genes [22], have a great influence on the organism development and represent the key elements in response to external and internal signals. Usually such genes reveal low to moderate transcriptional levels [23, 24] compared, for example, to genes of house-keeping machinery or structural genes. Still, such genes are actively transcriptionally regulated and assuming moderate absolute transcriptional levels, it may become difficult to differentiate between real changes in expression and random fluctuations. In this section we investigate if an accurate data cleaning step may assist the detection of such genes.

Here we are interested in detection of genes with low to moderate transcriptional, but high translational status, i.e. genes whose few mRNA copies intensively produce protein products. The criterion for the definition of such genes will be as follows:

- mRNA counts for gene *i* in total fraction is lower 300 (N_total,i_≤300, 7945 genes out of 23102);
- logarithm of the ratio of mRNA counts in polysomal and monosomal fractions is grater 1.5: *log_2_(N_polysomal,i_/N_monosomal,i_) ≥ 1.5*;
- significance (p-value) of the difference between polysomal and monosomal fractions identified by edgeR ≤ 10^−4^.

This criterion was applied to three datasets – raw data, data cleaned by setting a threshold for minimal accountable mRNA counts (24, 16 and 28 counts for total, monosomal and polysomal fractions respectively), and data cleaned by subtraction of the maximal “noise contributions” from the all mRNA counts (formula 1). The resulted gene lists were analyzed for functional annotation using DAVID [16] for the term “signal”. The keyword “signal” was selected, because it comprises genes involved in signaling pathways, like cytokines, gibberellin, auxin and ethylene signaling pathways regulating many aspects of plant growth and development including seed germination, stem and leafs, flower, pollen and fruit development *etc*. The results are presented in table 1.

**Table 1.**
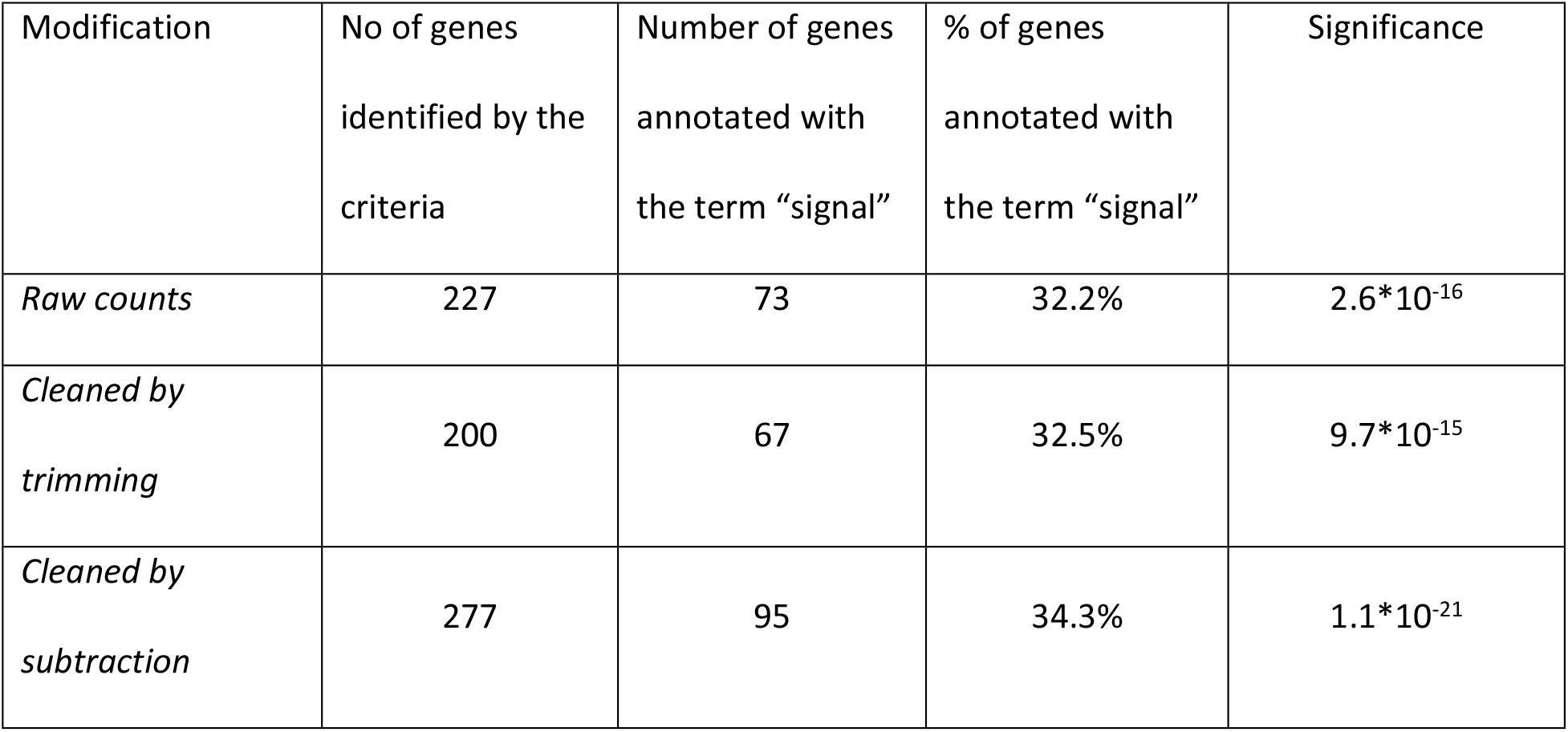
Genes with moderate to low transcription and high translation. Differentially translated genes were identified using EdgeR in three datasets: raw data, trimmed data and data cleaned by subtraction (see text for explanation). To limit the search to genes with moderate transcription, only genes with lower than 300 counts were considered (corresponds to approx. a lower third of all genes). Classification of genes using DAVID were performed to find genes with regulatory potential. Gene lists are available as supplementary material. Significance values as reported by DAVID.

It is evident from the table, that the data cleaning step is essential for detection of genes with regulatory function. The suggested cleaning via subtraction of the “noisy counts” results in detection of more genes, moreover, the percentage of regulation-related genes has also slightly increased. The results also support our hypothesis, that regulatory genes tend to show only moderate levels of transcription, but the most significant overrepresentation is observed for the data cleaned by subtraction (table 1).

Comparison of the identified gene sets revealed 122 genes found only using the data cleaned by subtraction, 72 genes found only by raw data and 155 genes found by both (gene lists are available in supplementary material). Focusing on genes annotated with “signal” term the corresponding numbers will be 39, 18, 56 (cleaned, raw and both datasets). This demonstrates, that the data cleaning procedure objectively extends the number of identified genes of interest. For example, there are such genes like root meristem growth factor (RGF3, AT2G04025), embryo-specific protein (ATS3, AT5G62210), transmembrane protein (DUF1191, AT4G23720) and many others directly related to gene regulation and signal transduction, all found exclusively after the suggested data cleaning.

It is interesting to note, that the commonly accepted approach to remove mRNA with counts below some pre-defined threshold leads to significantly fewer genes even compared to the raw data (table 1) and therefore, it was not used in the above comparisons. We also do not apply conventional pre-selected thresholds for the counts for the following reasons. First, the variation of those is quite significant and ranges from just a few in most studies [7, 9] to 32 counts [13] and the reasoning for preferring one to another is not evident. Second, even application of data-specific thresholds in the range of 16-28 led to significant reduction in number of identified genes, making this way of data cleaning ineffective. Programs like EdgeR or DESeq2 already have a built-in noise reduction logic, which probably makes the use of fixed thresholds unnecessary.

Another discussion point is the exponent estimation and how many data points should be included in more general cases. It can be suggested to use a local minimum in the area of small RNA counts as a last point. On the graph for total and monosomal fractions (fig. 2) this selection is quite evident. In contrast, data in polysomal mRNA fraction have greater variation, which objectively allows less exact estimation of parameters. Our investigation shows that as small as four points are sufficient to estimate the parameters of the exponent.

Overall, data modelling allows identifying characteristics of exponential distribution and thereby to exclude possible noise from the measured mRNA counts. Such data modification allows to fine-tune the conventional search algorithms, especially when genes with moderate transcriptional levels are in focus.

### 3.4. Detailed functional analysis

The use of functional classification of genes like Gene Ontology is practical to give a quick overview on underlying differences in functionality of the investigated genes. Here the resource PANTHER v.14.0 [17] was used to classify the mRNAs in four datasets. These datasets were compiled using “symmetrical” criteria to the criterion defined above. Particularly, mRNA are classified according to the level of transcription into low and high (*N_total,i_*≤300 and *N_total,i_*≥1200, respectively) and according to the level of translation into monosomal and polysomal mRNAs (*log_2_(N_polysomal,i_/N_monosomal,i_)* ≤ −1.5 and ≥ 1.5 respectively, in both cases p-value by edgeR ≤ 10^−4^). The values of 300 and 1200 for total mRNA were selected as the lowest and highest 3-quantiles of all genes (7945 and 7846 genes respectively). The four datasets comprise 330, 444, 277 and 473 genes (high & polysomal, high & monosomal, low & polysomal and low & monosomal respectively) and are available in the supplementary material.

PANTHER classification system is designed to classify genes according to families of evolutionary related proteins, protein molecular functions, pathways etc. The four datasets were classified according to Gene Onthology (GO) molecular function and PANTHER protein class categories, the latter is used to categorize protein families (fig. 3). Classification by GO “molecular function” demonstrate the significant overrepresentation of genes with molecular function “regulator” (GO:0098772) in the polysomal mRNAs with low transcription (p-value=5.89*10^−5^, observed 14.5%, expected 3.2%, here and further binomial test, fig. 3A dark blue slice marked with *). Genes in this category include, for example, cyclin-B1, root meristem growth factors, pectinesterase inhibitors. Corresponding category in PANTHER protein class “gene specific translational regulator” (PC00264) is also overrepresented only in the same mRNA group (p-value=2.07*10^−4^, observed 11.6%, expected 2.0%, fig. 3B). To regulator-related could also be regarded genes with a function of molecular transducers (GO:0060089, p-value= 1.87*10^−3^, observed 7.3%, expected 1.6%), which work as compound molecules with one or more regulatory components. Genes involved in pore formation regulating the transit of other of molecules (transporter activities) are also overrepresented in low transcribed genes (p-value=2.64*10^−6^, observed 10.9%, expected 2.4%) with no preference to polysomal or monosomal mRNA groups. This particularly may indicate potential active differential regulation of translation of genes in this group.

**Figure 3.**
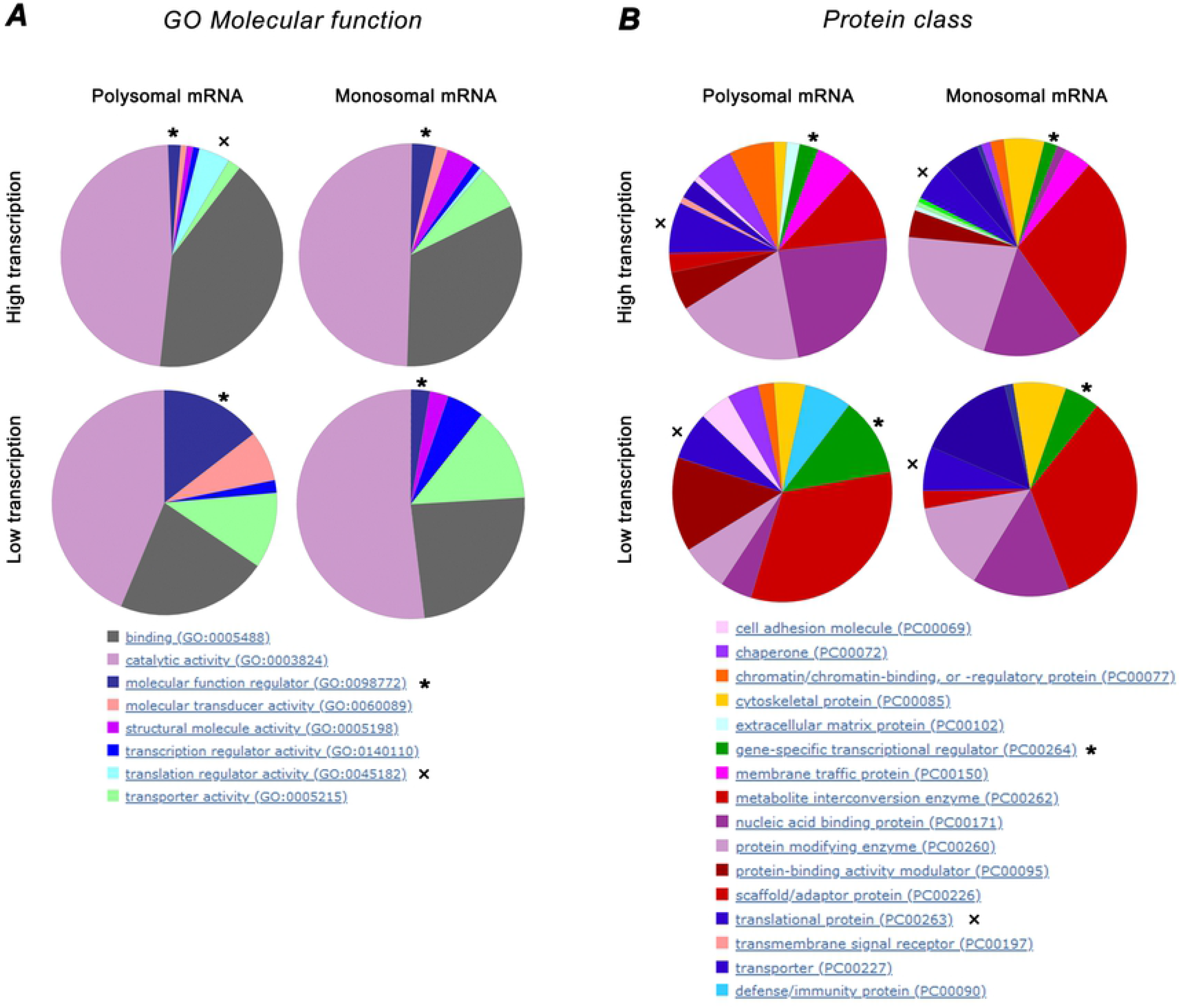
Functional classification of mRNA depending on transcriptional and translational status. mRNAs were classified into four groups according to transcriptional and translational levels (see text). A. Classification using GO “molecular function” demonstrates the significant overrepresentation of genes with molecular function “regulator” in the mRNA with low transcription and high translation (p-value=5.89*10-5, dark blue slice marked with *). Regulation related “translational regulator” group shows only moderate significance (p-value= 8.38*10-3, marked with x) in the group of genes with high transcription. B. Classification according to “protein class” by PANTHER classification system. Similarly, transcriptional regulator genes are significantly overrepresented (p-value=2.07*10-4, green slice marked with *). Translational proteins do not reveal any significant biases (dark blue slice marked with x).

An interesting exception is the group of “translational regulators” (GO:0045182), which is represented only in highly transcribed genes, although the significance is only at the moderate level (p-value=8.38*10^−3^, observed 4.6%, expected 1.6%, fig. 3A marked with x). Genes classified into this group are genes of a close family of eukaryotic translation initiation factors: eIF-2, 4B2, 4B3, 4G and Ts. Therefore, we may speculate, that high transcription of the above translation initiation factors cannot be extrapolated on all genes related to regulation of translation, because it is not confirmed by the “protein class” classification scheme, by which translation related genes are equally distributed among groups (PC00263, fig. 3B marked with x). The above genes may represent a closely related gene family with similar transcriptional regulation, that may indeed have high transcriptional levels and is an exception to the general rule, or it could be just a statistical artefact.

## Conclusion

Investigation of regulatory genes is crucial for the understanding of the functioning of any organism, but the experimental detection of such genes is complicated by the low to moderate levels of their expression and the significant influence of experimental and biological noise. One way to overcome this is to investigate target genes with strong expression and apply reverse engineering or use databases of regulatory pathways to find the regulators. Direct methods utilize complex mathematical models to discern weak signals of regulation.

The data cleaning procedure suggested here is assumed not to further complexify the methods, but to “personalize” parameters, used to dissect noise and real values. The idea consists in defining a maximum contribution, which could originate from technical or biological noise, with a subsequent subtraction of that value from the raw measurements. This is different to other approaches, where only values below some noise threshold are removed and the rest is left intact. As shown in the results, the suggested cleaning procedure increases the number of detected genes with differential expression. Moreover, the ratio of genes with regulatory functions is also increased after suggested data cleaning.

We believe that data modelling should be used to define dataset–specific thresholds and the use of “universal” values avoided, since variation caused by experimental settings could be significant. The polysomal and monosomal fractions in our experiment differs almost twice in the level of the introduced noise, despite standardized sample preparation and sequencing procedures. The suggested in the literature threshold values cover a very broad range, so the selection of a particular threshold to our view needs transparent justification, no matter if they are used to trim the low values or to clean the data as suggested here.

Finally, the suggested experimental design to measure three mRNA fractions allows investigation of both quiet and highly translated mRNA, since the investigation of potential mechanisms of translational repression are of the same importance as mechanisms of activation. Understanding of both will provide the complete picture of translational regulation.

## Acknowledgments

We are grateful Dr. Charles Latting for thorough English editing.

## Supporting information

Excel file with gene lists

